# Developmental Plasticity and Stromal Co-option Shape a Pituitary Neuroendocrine Tumor Transcriptional Continuum

**DOI:** 10.1101/2025.06.18.659743

**Authors:** Robert C. Osorio, Jun Y. Oh, Ankita Sati, Jangham Jung, Alexander J. Ehrenberg, Atul Saha, Meeki Lad, Harmon Khela, Nicole Brennick, Petros Giannikopoulos, William W. Seeley, Lea T. Grinberg, Aaron Diaz, Manish K. Aghi

**Affiliations:** Department of Neurological Surgery, University of California San Francisco, San Francisco, CA; Innovative Genomics Institute, Berkeley, CA; Department of Neurology, Weill Institute of Neuroscience, Memory and Aging Center, University of California San Francisco, San Francisco, CA; Department of Laboratory Medicine, University of California, San Francisco, CA

**Keywords:** pituitary adenoma, pituitary tumor, PitNET, tumor microenvironment, single-nuclei RNA sequencing, spatial transcriptomics, prolactinoma, Cushing’s

## Abstract

Pituitary neuroendocrine tumors (PitNETs) are common intracranial neoplasms with complex biology underpinned by unresolved cellular origins, molecular heterogeneity, and microenvironment interactions. Here, we employ single-nuclei RNA-sequencing (snRNA-seq) of 419,874 cells from human normal pituitaries and PitNETs with spatial transcriptomics to resolve these challenges. We identify multi-hormonal neuroendocrine cells in both normal and tumor tissues, originating as early pseudotime intermediates from pituitary stem cells, revealing an inherent plasticity that blurs traditional lineage boundaries. PitNETs exhibit a transcriptional continuum across subtypes, challenging their classification into discrete categories. Trajectory analysis uncovers divergent cellular origins: silent gonadotroph adenomas (SGAs), prolactinomas, silent corticotroph adenomas (SCAs), and Cushing’s adenomas are closely linked to differentiated neuroendocrine cells, while somatotroph and null cell adenomas (NCAs) appear to derive more directly from adult stem cells. Tumor cells co-opt robust cell-cell communication networks found in normal adult neuroendocrine cells. Spatial profiling further demonstrates that perivascular niches enhance tumorigenicity through angiogenic and epithelial-mesenchymal transition programs. Our work redefines PitNETs as ecosystems shaped by developmental plasticity and microenvironmental crosstalk, offering a roadmap for future therapies targeting lineage fluidity and stromal dependencies.

## 1. Introduction

Pituitary neuroendocrine tumors (PitNETs, previously known as pituitary adenomas) are common intracranial tumors originating from the anterior pituitary gland. Nearly half of PitNETs overproduce hormones and cause symptoms specific to their neuroendocrine type. For instance, hypersecreting lactotroph tumors (prolactinomas) lead to galactorrhea. Hypersecreting somatotroph tumors cause acromegaly, while hypersecreting corticotroph tumors lead to Cushing’s disease. In contrast, some PitNETs remain hormonally silent and primarily manifest as mass effects or are incidentally diagnosed. While most silent PitNETs are silent gonadotrophs, all types of hypersecreting PitNETs can have a silent counterpart.

The 2017 World Health Organization (WHO) classification brought a paradigm shift by categorizing pituitary tumors based on cell lineage, determined through the immunohistochemical expression of three transcription factors (TFs) implicated in the development of the various subtypes of normal neuroendocrine cells, with TPIT (TBX19) immunopositivity identifying corticotrophic PitNETs; PIT1 immunopositivity identifying somatotroph, lactotroph, or thyrotroph PitNETs; SF1 immunopositivity identifying gonadotroph PitNETs, and triple-negative immunostaining identifying null cell PitNETs. This new approach deviates from the previous reliance solely on hormone type and is maintained in the 2022 WHO classification^1,2^. While the pituitary cell lineage classification system represents a notable insight, the complex process of PitNET tumorigenesis and the dynamics of TF expression are not yet fully understood. Additionally, it remains uncertain whether the presence of PitNETs with multiple transcriptional signatures or varying degrees of differentiation^3,4^ necessitates the need for a more flexible subtyping approach, as their nature may be more fluid than previously presumed.

Previous molecular studies of PitNETs have been constrained by methodological limitations. Bulk RNA sequencing obscures cellular diversity^4^, while prior single-cell analyses^3,5–7^ hindered by small cohorts, cell dissociation artifacts, and sparse spatial resolution—fail to resolve rare subpopulations or microenvironmental interactions. Here, we leverage snRNA-seq to profile 419,874 nuclei from 11 PitNETs and two normal pituitary glands, bypassing dissociation bias in fragile neuroendocrine cells^8,9^, and integrate spatial transcriptomics. We map the developmental trajectories of PitNET subtypes, decode their stromal crosstalk, and reconstruct a high-resolution atlas of human pituitary glands and PitNETs. Our findings redefine PitNETs as ecosystems shaped by lineage fluidity and microenvironmental hijacking, offering a roadmap to refine diagnostic and therapeutic frameworks.

## 2. Results

### 2.1 Single-nuclei atlas of adult human pituitary glands

To elucidate the cellular composition and transcriptional dynamics of the normal adult human pituitary gland, we performed single nuclear RNA-sequencing (snRNA-seq) on postmortem pituitary glands collected from non-tumor subjects (n=2) with short postmortem intervals (PMI < 10 hours) **(Fig. 1a).** After implementing rigorous quality control and filtering steps, we obtained 29,071 high-quality nuclei, with an average of 6,540 unique molecular identifiers (UMIs) and 2,643 genes per nucleus. To reduce the dimensionality and identify nuclei with similar transcriptomic patterns, we implemented Principles Component Analysis (PCA) and Uniform Manifold Approximation and Projection (UMAP) algorithm using Seurat software^10^, resulting in 17 distinct nuclei clusters present in both samples **(Fig. 1b-d)**. Using canonical gene markers from previous studies^3,11^ and SingleR^12^, we identified all five types of neuroendocrine cells (corticotroph, gonadotroph, lactotroph, thyrotroph, somatotroph), immune cells (macrophage, T cells), stromal cells (pituicyte, pericyte, endothelial cells), and adult pituitary stem cells **(Fig. 1b, e)**. Each cluster contained cells from both donors, and the percentage of cells from each sex was similar, suggesting minimal batch effect **(Fig. 1c, d)**.

**Figure 1:**
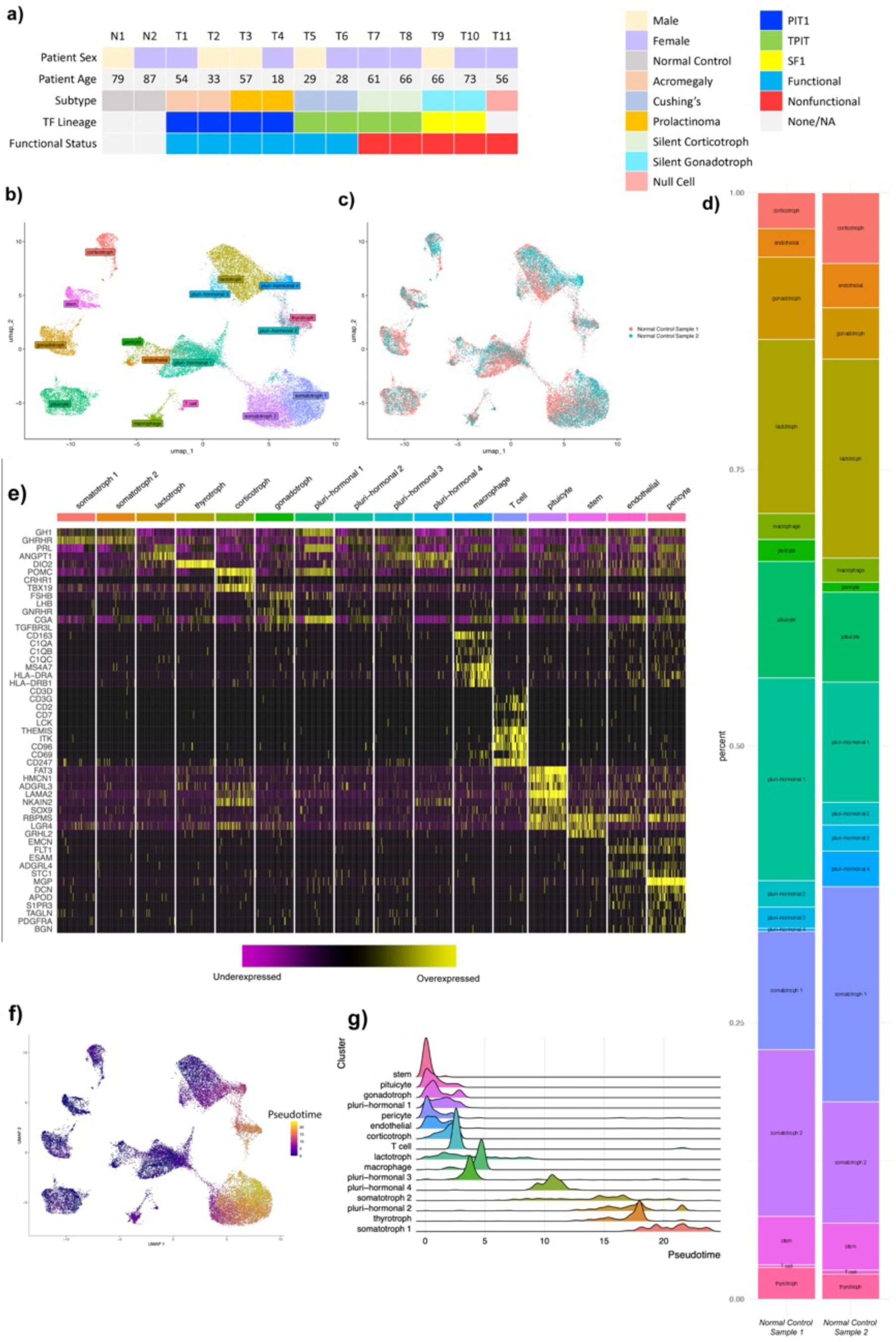
Study overview and single-nucleus transcriptomic atlas of the normal human pituitary. (**a**) Overview of study cohort and tissue samples. (**b**) Dimensional reduction plot of human pituitary gland nuclei, clustered by cell identity. (**c**) Dimensional reduction plot of human pituitary gland nuclei, clustered by patient origin of tissue sample. (**d**) Stacked bar chart of relative cluster sizes in each normal control sample. (**e**) Transcriptomic heatmap of RNA expression of canonical markers expressed in the adult pituitary gland. (**f**) Dimensional reduction plot of human pituitary gland nuclei after trajectory analysis, colored by pseudotime. (**g**) Ridge plot of normal pituitary clusters demonstrating pseudotime distribution of each cluster and ordered by mean pseudotime. TF: transcription factor.

Consistent with previous findings on pluripotent stem cells in the anterior pituitary^13–16^, our stem cell cluster exhibited high expression of genes *RBPMS*, *LGR4*, and *GRHL2* **(Fig. 1e)**. The stem cell cluster also expressed a greater number of TF genes than mature neuroendocrine cells, including TF genes previously described in human fetal pituitary (i.e., *PROP1*, *NFIB*, *LHX3*, *HEY1*, *ZNF521*)^11^ and 48 novel pituitary TF genes **(Supplementary Table 1).** Gene ontology analysis of the upregulated genes in the stem cells revealed enrichment in processes related to epithelium development and cell morphogenesis, consistent with their stem-like identity.

An interesting finding emerging from our single nuclei atlas of the adult human pituitary gland was four clusters (20% of total cells in normal control samples) exhibiting co-expression of various neuroendocrine transcriptomic signatures. These clusters were designated pluri-hormonal clusters 1-4 as follows: (1) pan-hormonal (somatotroph/lactotroph/corticotroph/gonadotroph), (2) somatotroph/gonadotroph, (3) somatotroph/lactotroph, also known as somatomammotroph,^17^ and (4) lactotroph/thyrotroph **(Fig. 1b, e)**. Previous studies using immunohistochemistry have observed PIT-1 lineage plurihormonal tumor cells^2,18,19^, rarer multi-lineage plurihormonal tumor cells^20^, and the presence of plurihormonal cells in the normal adult human anterior pituitary.^21^ However, the classification of plurihormonal cells as distinct cell lineage or as intermediate stages between different lineages remains a subject of debate.^22–24^ To investigate further, we conducted pseudotime trajectory analysis^25,26^ to explore the developmental progression of plurihormonal cells relative to stem cells and classical monohormonal neuroendocrine cells **(Fig. 1f, g)**. We found that monohormonal gonadotroph and corticotroph cells diverged from stem cells early in our pseudotime, suggesting early differentiation of TPIT and SF-1 lineage, respectively, with few intermediate states. This finding differs with a previous snRNA-seq study involving 4113 cells, which reported more intermediate states for both gonadotroph and corticotroph cells.^11^ In contrast to TPIT and SF-1 lineage, PIT1-lineage cells (i.e., lactotroph, somatotroph, and thyrotroph cells) emerged in the middle or late stages of pseudotime **(Fig. 1g).** Interestingly, pan-hormonal cluster appeared early in pseudotime, whereas plurihormonal clusters 2, 3, and 4 emerged later **(Fig. 1g)**, suggesting that, while the latter plurihormonal clusters represent terminal states with possible secreting functional capability, pan-hormonal cluster, which co-expresses transcription markers for four endocrine type cells, might represent a transient, intermediate state that undergoes additional pruning or specialization in response to physiological demands or pathological stimuli. ^27–29^

Next, we leveraged pseudotime analysis to identify key TFs associated with the development of each endocrine cell type from stem cells **(Supplementary Figs. 1-8; Supplementary Table 2)**. Certain TF genes, such as *ST18* and *ZBTB16*, exhibited increased expression across multiple cell lineages, suggesting their broader role in general neuroendocrine differentiation Notably, *MYT1L* and *ZNF10* displayed increased expression in all three PIT-lineage neuroendocrine cells, while *NR4A1* exhibited specific upregulation in thyrotroph and lactotroph cells. Estrogen Receptor 1 (*ESR1*), a ligand-activated TF, was associated with both gonadotroph and lactotroph maturation, in line with immunohistochemistry studies demonstrating high expression of estrogen receptors in PRL, FSH, or LH-positive endocrine cells^30^. Additionally, nuclear receptor 4A2 (*NR4A2*), a member of the steroid-thyroid hormone-retinoid receptor superfamily which directly binds the *PRL* promoter^31^ showed increased expression during lactotroph development **(Supplementary Fig. 4).** *NR4A2* is implicated in the establishment of dopaminergic systems in the brain without a canonical ligand-binding domain, featuring instead a narrow and tight cavity for small molecular ligands such as prostaglandins to bind^32,33^.

Furthermore, cells in the four plurihormonal clusters we had identified expressed TF genes that generally mirrored their hormonal-combination phenotype. For instance, in plurihormonal cluster 3 (somatotroph/lactotroph), we observed upregulation of somatotrophic differentiation-associated TF genes (i.e., *HIF3A*, *PLAGL1*, *ZBTB16*) and lactotrophic differentiation-associated TF genes (i.e., *ESR1*, *NR4A1*, *NR4A2*) over pseudotime, suggesting additive effects of TF activities in these plurihormonal cells. Finally, we found that many stem cell-associated TF genes, including previously described genes (i.e., *GRHL2*, *HMGA2*, *MECOM*, *NFIB*, *TCF7L1*, *ZNF521*^11^) and other novel genes (i.e., *RFX4*, *BNC2*, *EBF1*, *NFIB*, *SETBP1*, *NPAS3*, *NFIA*) were downregulated during development across multiple cell lineages (**Supplementary Fig. 1-8)**.

### 2.2 Single-nuclei characterization of PitNET subtypes

We then collected samples from 11 PitNETs (acromegaly (n=2), Cushing’s adenoma (n=2), silent corticotroph adenomas (SCA) (n=2), prolactinoma (n=2), silent gonadotroph adenomas (SGA) (n=2), and null cell adenoma (NCA) (n=1)) during transsphenoidal surgeries for snRNA-seq. All tumor samples underwent immunohistopathological examination of canonical PitNET markers using standard criteria^2^ **(Fig. 1a)**. After quality control, we obtained 390,718 nuclei from tumor samples, with an average of 327 UMIs and 116 genes per nucleus. We then reconstructed UMAP with 390,718 nuclei from tumor samples and 29,071 nuclei from control samples **(Fig. 2a, b)**, yielding 29 distinct clusters.

**Figure 2:**
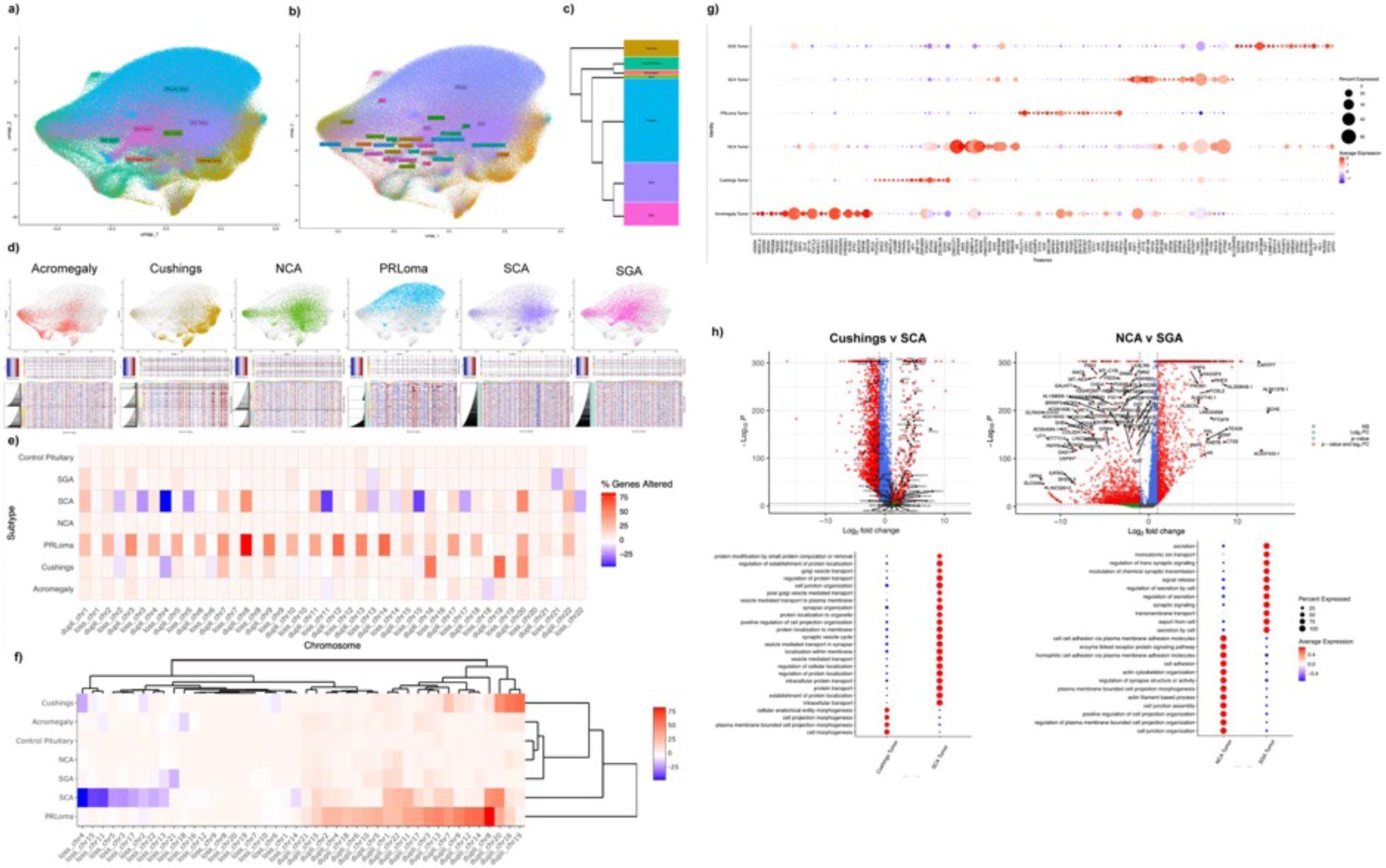
Transcriptomic and copy number variation (CNV) analysis of pituitary neuroendocrine tumors. (**a**) Dimensional reduction plot of pituitary neuroendocrine tumors and nontumor nuclei from all samples, integrated and clustered by tumor subtype vs nontumor status. (**b**) Dimensional reduction plot of pituitary neuroendocrine tumors and nontumor nuclei from all samples, further subclustered to display the identities of nontumor nuclei. (**c**) Clustering of nuclei based on tumor subtype and nontumor status, based on overall pseudobulk transcriptomic expression. (**d**) Relative contributions and locations of each tumor subtype to the overall dimensional reduction (upper), followed by CNV analysis of each tumor subtype relative to nontumor controls (lower). (**e**) Heatmap of aggregated CNV data, plotting relative percent of genes altered per chromosome and split by tumor subtype vs normal control. Ordered by chromosome number and CNV gain vs loss. (**f**) Heatmap of aggregated CNV data, clustered by similar CNV expression. (**g**) Transcription factor analysis of differentially expressed genes (DEGs) of each tumor subtype. (**h**) Volcano plots of DEGs (upper) and gene ontology pathways (lower) differentially expressed between Cushings and silent corticotroph adenomas (left) as well as between null cell adenomas and silent gonadotroph adenomas (right). NCA: null cell adenoma; PRLoma: prolactinoma; SCA: silent corticotroph adenoma; SGA: silent gonadotroph adenoma; CNV = copy number variant.

At the snRNA transcriptomic level, integration of PitNET samples revealed distinct tumor clusters with significant overlap, indicating a continuum within the subtypes rather than strict classification (**Fig. 2a, b)**. Hierarchical clustering based on the entire transcriptome further showed that silent corticotrophic PitNETs clustered closely with silent gonadotroph PitNETs, while being distant from secreting corticotrophic PitNETs **(Fig. 2c)**, suggesting a greater shared transcriptome across silent PitNETs than across PitNETs with a shared hormonal staining but variable secretion patterns. Furthermore, despite belonging to the PIT1 lineage family, prolactinomas and acromegaly adenomas did not cluster together **(Fig. 2c).** These findings supported previous speculation^4,34^ that lineage-specific TPIT, SF1, and PIT1 classification alone may not accurately reflect the cellular identities of these tumors.

We next examined large-scale chromosomal instability in pituitary tumors using single-cell large-scale copy-number variations (CNVs) inferred by snRNA-seq **(Fig. 2d)**. Previous studies suggest that, despite their predominantly benign nature, PitNETs often display arm-level copy-number alterations across substantial portions of the genome^35^, and these disruptions are present in the majority of individual tumor cells^5^. Compared to control cells from the normal pituitary gland, we observed the most increased CNVs across the genome in prolactinomas, while immune-negative null-cell adenoma showed the least alterations in CNV **(Fig. 2d, e),** providing single-cell corroboration of previously reported microarray findings^36^. Unsupervised hierarchical clustering of PitNETs based on CNV profiles revealed two key findings. First, CNV changes are specific to tumor subtypes, with somatotrophic and null cell PitNETs showing the least CNV changes and lactotrophic PitNETs showing the most CNV changes (**Fig. 2f**). Second, silent corticotrophic PitNETs are a distinct subtype with a unique CNV profile far more altered than secreting corticotrophic PitNETs rather than a subtle modification of secreting corticotrophic PitNETs **(Fig. 2f).**

Our snRNA-seq transcriptome-wide analysis corroborated previously described gene expression biomarkers^3,37^ for somatotrophic (e.g. *GH1*, *GHRHR*, *ENPP1*), secreting corticotrophic (e.g. *POMC*), silent corticotrophic (e.g. *TBX19*), lactotrophic (e.g. *PRL*, *ENO3*, *NTS*), and silent gonadotrophic (i.e., FSHB, CHGA, TGFBR3L) PitNETs. Null-cell adenomas were negative for these specific genes. We discovered novel TFs that are tumor subtype-specific, including somatotrophic (*HOB4*, *NKX2-2*, *HOB3*, *NPAS4*, *PRDM8*), secreting corticotrophic (*TFCP2L1*, *BNC2*, *CUX2*, *ARNTL2*, *LIN28B*), lactotrophic (*VDR*, *ZNF831*, *ZNF521*, *RARB*, *CXXC5*), silent gonadotrophic (*TCF7*, *L3MBTL3*, *ZNF213*, *SCRT1*, *PROX1*), and NCA (*ONECUT1*, *ZNF804A*, *MKX*, *PRRX1*, *L3MBTL4*) **(Supplementary Table 3)**. Notably, we also identified increased expression of TF genes, such as *PPARG*, *NR2F2*, and *SOX5*, in PitNETs that can be targeted using ligand inhibitors (e.g. thiazolidinedione) or miRNAs^38–42^ **(Fig. 2g)**. Notably, somatotrophic PitNETs from acromegalic patients highly expressed *CEBPD* **(Fig. 2g),** which actively represses other PIT1 lineage differentiations during normal development^11,43^, suggesting that PitNETs may hijack inhibitory mechanisms to dictate and maintain a specific hormonal phenotype. Additionally, we noted upregulation of specific TF genes (e.g. *YBX3*, *SREBF1*, *ETV5*, *NPAS2*, and *TRPS1*) that have been implicated in the pathogenesis of other neuroendocrine and non-neuroendocrine tumors, including colorectal, nasopharyngeal, prostate, and breast cancers. **(Fig. 2g)**^44–48^. These findings lend support to the controversial 2022 WHO classification of pituitary tumors as encompassing features of both neuroendocrine and non-neuroendocrine tumors ^49^.

We next compared the transcriptomes of secreting versus silent corticotrophic PitNETs given their presumed origin from a common TPIT lineage cell. As expected, TPIT was equally present in both secreting and silent corticotrophic PitNETs, and secreting corticotrophic PitNETs showed increased expression of POMC **(Fig. 2h)**, a key precursor of ACTH absent in silent corticotrophic PitNETs. Surprisingly, and consistent with our CNV findings, there were far more differences between secreting versus silent corticotrophic PitNETs than just POMC loss in the latter, as we found secreting corticotrophic PitNETs to be less mature and differentiated than silent corticotrophic PitNETs, with secreting corticotrophic PitNETs displaying higher expression of genes involved in morphogenesis and cellular development compared to silent corticotrophic PitNETs **(Fig. 2h)** and silent corticotrophic PitNETs expressing genes involved in intracellular vesicular transport, localization, and synaptic release, indicating retained functional-structural potential characteristic of a mature state **(Fig. 2h).** Silent corticotrophic PitNETs also had higher expression of *PCSK1N*, an indirect inhibitor of ACTH maturation **(Fig. 2h)**, potentially actively repressing hormone production in ways beyond their previously demonstrated loss of POMC expression.

The classification of NCAs and SGAs as distinct subgroups of nonfunctioning PitNETs is still a subject of debate^37,50^. Our analysis reveals that despite being non-functioning, SGAs exhibit gene expression patterns related to hormonal secretion, similar to SCAs. **(Fig. 2h)**. In contrast, NCAs exhibit upregulation of genes involved in cell-cell adhesion and communication, indicating the presence of a robust intercellular network akin to that found in normal neuroendocrine cells **(Fig. 2h).**

### 2.3 Cellular trajectories of PitNET subtypes

The cellular origin of PitNETs remains uncertain,^51–53^ with two prevailing hypotheses. The first hypothesis is that PitNETs originate from the oncogenic transformation of cells in the normal pituitary gland with a stem-like transcriptome that we identified in our analysis of normal adult pituitary glands above **(Fig. 1b, e)**. The second hypothesis is that PitNETs derive from differentiated neuroendocrine cells in the normal gland that undergo clonal de-differentiation to produce a tumor-initiating stem-like cell with an endocrine phenotype.^54^ To identify the proximate precursor cells of each PitNET subtype, we conducted pseudotime trajectory analysis using adult pituitary stem cells in the normal pituitary gland, mature neuroendocrine cells in the normal pituitary gland, and PitNET cells **(Supplementary Table 4).** Interestingly, we found that PitNETs exhibited both origin patterns depending on their subtype **(Fig. 3a-e)**.

**Figure 3:**
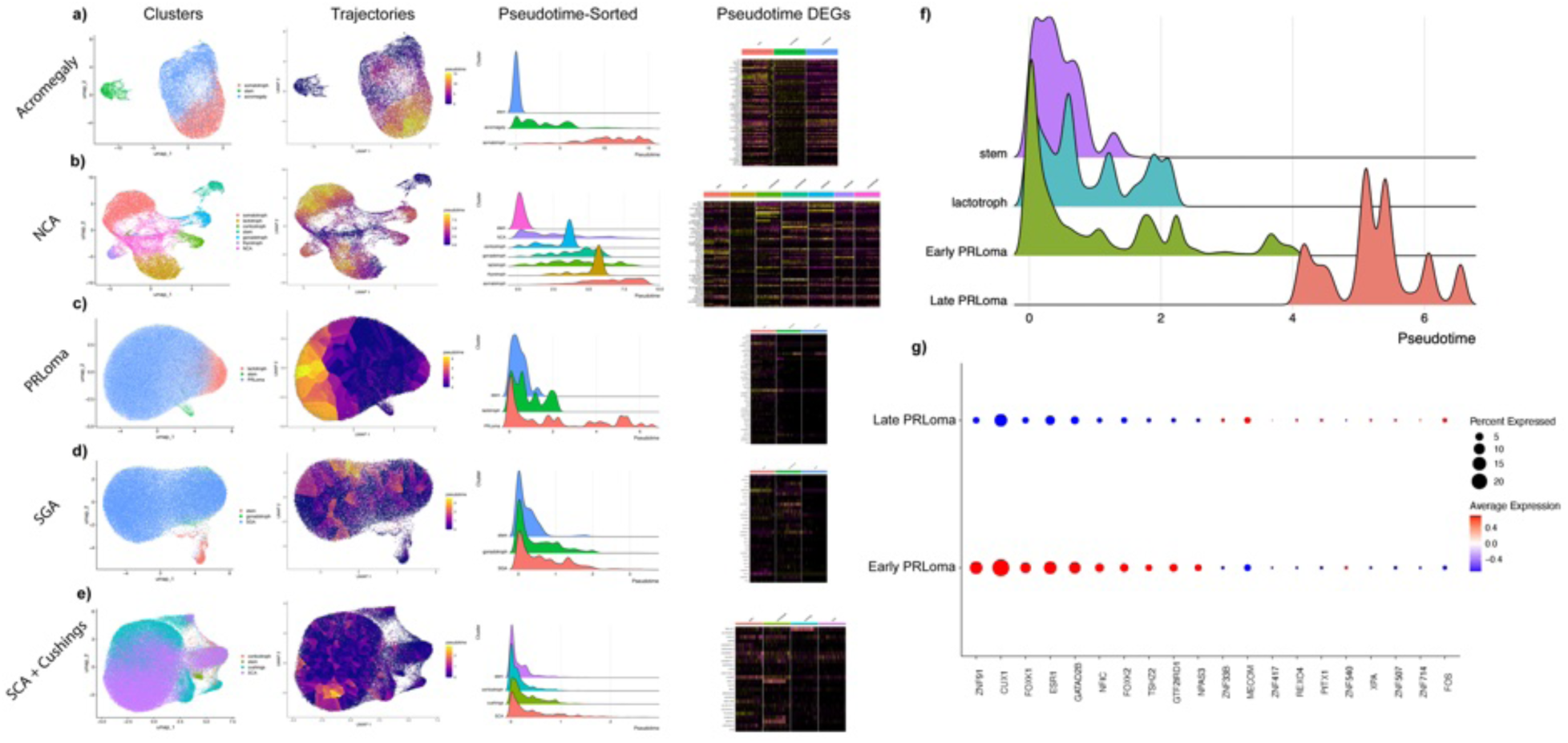
Pseudotime trajectory analysis of pituitary neuroendocrine tumors. (**a-e**) Dimensional reduction of each tumor subtype, plotted with stem, tumor nuclei, and normal neuroendocrine counterpart nuclei. For null cell adenomas, all neuroendocrine nuclei were included (“Clusters”). Similar plots re-colored by pseudotime after trajectory analysis (“Trajectories”). ridge plots of demonstrating pseudotime distribution of each cluster and ordered by mean pseudotime (“Pseudotime-Sorted”). Transcriptomic heatmaps of RNA expression of pseudotime-correlated differentially expressed genes with either increased or decreased expression in each cluster (“Pseudotime DEGs”). (**f**) post-hoc ridge plot of prolactinoma nuclei after splitting prolactinomas into early and late subclusters. (**g**) pseudotime-correlated DEGs of early and late prolactinoma nuclei. NCA: null cell adenoma; PRLoma: prolactinoma; SCA: silent corticotroph adenoma; SGA: silent gonadotroph adenoma; DEG: differentially expressed gene.

Normal adult gonadotroph cells were intermediaries between normal stem cells and SGA tumor cells, indicating the likely origin of SGAs from differentiated gonadotroph cells **(Fig. 3d)**. Similarly, all lactotrophic PitNET cells emerged after normal mature lactotroph cells in the trajectory and were further categorized into early-phase (Early-Prl) and late-phase (Late-Prl) **(Fig. 3c, f).** Early-Prl cells highly expressed TF genes such as *ZNF91*, *CUX1*, *FOXK1*, *ESR1*, and *GATAD2B*, whereas Late-PRL cells predominantly expressed *ZNF33B*, *MECOM*, *PITX1*, and *FOS* **(Fig. 3f, g)**. We also examined the pseudotime ordering of Cushing’s adenomas and SCA, based on our transcriptomic data suggesting a more differentiated phenotype for SCA **(Fig. 2e)**. Both tumor types emerged after normal mature corticotroph cells, and SCA emerged at a later stage than Cushing’s adenomas **(Figure 3e),** supporting our earlier findings **(Fig. 2h)**.

In contrast to these PitNET subtypes which appeared to arise from differentiated neuroendocrine cells in the normal gland, somatrophic PitNET cells appear early in the trajectory from stem cells, prior to normal mature somatotroph cells **(Fig. 3a)**, suggesting a close link with glandular stem cells rather than mature endocrine cells. NCAs, which lack differentiation markers as indicated by IHC and specific lineage-specific TFs, remain enigmatic in their origins, though they are occasionally speculated to be gonadotrophic.^55^ Our reconstructed pseudotime trajectory incorporating NCAs, stem cells, and all five types of mature neuroendocrine cells shows that NCAs emerge early, immediately following normal stem cells **(Fig. 3b)**, suggesting that their origins are more closely linked to normal stem cells, rather than any mature endocrine cell types. Interestingly, both of these subtypes that appeared to originate from stem cells in the normal pituitary gland (somatotrophic PitNETs and NCAs) also had the least CNV changes, a finding that contrasts malignant cancers where cancers originating from cells with greater plasticity tend to be associated with more CNV changes.^56,57^

### 2.4 Cell-to-cell communication networks in the normal pituitary gland and in PitNETs

Previous studies have speculated that the anterior lobe of the pituitary gland is not merely a patchwork of hormone-secreting cells but rather a well-organized network capable of close-distance crosstalk and co-oscillation, enabling a coordinated response to physiological stimuli^58–60^. Using CellChat^61^, we first identified 46 diverse cell-cell signaling pathways in the normal anterior lobe of the pituitary gland based on ligand-receptor expressions. These pathways include NRG, NRXN, CADM, NCAM, VISFATIN, ALCAM, PDGF, and VEGF signaling **(Fig. 4a, b; Supplementary Table 5).**

**Figure 4:**
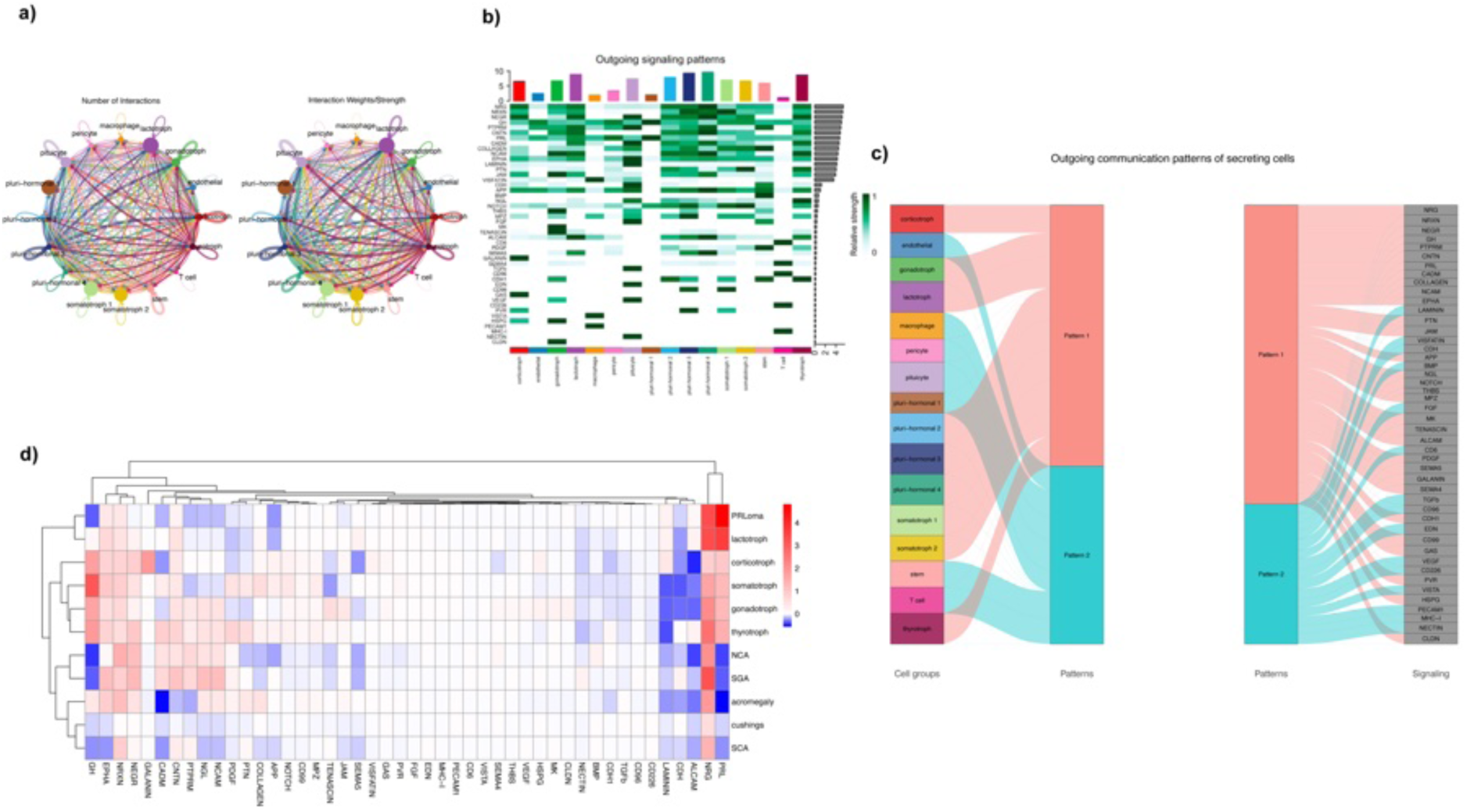
Cell-cell communication network analysis of pituitary neuroendocrine tumors. (**a**) overall communication size and strength between nuclei types. (**b**) expression of outgoing signaling ligands of nontumor nuclei in normal control samples. (**c**) expression of outgoing signaling ligands of tumor nuclei and their normal neuroendocrine counterpart nuclei, clustered by expression similarity. (**d**) river plot meta-clustering of normal control nuclei, where normal nuclei are sorted into “patterns” based on outgoing expression of communication ligands (left). Legend demonstrating the signaling ligands associated with each “pattern” (right). NCA: null cell adenoma; PRLoma: prolactinoma; SCA: silent corticotroph adenoma; SGA: silent gonadotroph adenoma.

Subsequent unsupervised clustering revealed two distinct cell-cell communication patterns in the normal anterior lobe of the pituitary gland: Pattern A, characterized by neuroendocrine cells communicating with different subtypes of neuroendocrine cells via pathways associated with cell-cell adhesions and junctions, suggesting their engagement in close communication networks and crosstalk^59,62^; and Pattern B, consisting of non-neuroendocrine cells communicating with pituitary stem cells through pathways related to immune, vascular, and other cellular functions **(Fig. 4c)**. Notably, within Pattern A, we identified the ephrin-ephrin receptor signaling pathway, a subset of receptor tyrosine kinase capable of bidirectional intercellular signaling. This pathway has been shown to play a crucial role in facilitating communication between β cells of the pancreatic islets for glucose-insulin homoestasis^63,64^, suggesting potential utilization of this network broadly across organs within the endocrine system engaging in physiological homeostasis. Interestingly, pan-hormonal cluster, emerging early in pseudotime **(Fig. 1d)**, lacked the typical pattern A cell-cell communication characteristics seen in other neuroendocrine cell types and clustered closer to the pattern B communication pattern seen with pituitary stem cells (**Fig. 4c)**, further indicating its incompletely differentiated and highly plastic state.

We next examined if PitNET tumor cells retain these communication pathways we had identified for cells in the normal pituitary gland **(Fig. 4d).** Surprisingly, PitNET cells closely mirrored normal neuroendocrine cell signaling, with high expression of pathways like NRG, NRXN, NEGR, and CNTN. EPHA was also highly expressed in pituitary tumor cells, except in Cushing’s and SCA cells **(Fig. 4d).** These findings suggest that PitNET cells utilize many examples of the two patterns of cell-to-cell communication we had identified in normal pituitary cells, and we speculate that close communication networks^65^ may be crucial for tumor cell survival and growth.

### 2.5 PitNET microenvironment

The tumor microenvironment is a highly structured ecosystem, consisting of tumor cells surrounded by diverse cell types and often embedded within an altered, vascularized extracellular matrix^6,66,67^. Our PitNET samples predominantly consisted of tumor cells, but we also detected various non-tumor cells, including normal neuroendocrine cells (corticotrophs, gonadotrophs, lactotrophs, and somatotrophs), endothelial cells, pituicytes, macrophages, mixed stroma cells, and stem cells. Plurihormonal cells, including pan-hormonal (i.e., cells with a somatotroph/lactotroph/corticotroph/gonadotroph signature), were also present in the PitNETs. We first characterized TF and non-TF DEGs in the non-tumor cells **(Fig. 5a)** and then examined subtle transcriptomic differences between non-tumor cells in PitNET samples and matched normal cell types in control samples **(Fig. 5b-l; Supplementary Table 6)**. Overall, normal neuroendocrine cells in PitNETs had more downregulated genes compared to matched cell types in control samples, except in lactotroph PitNETs, whose normal neuroendocrine cells exhibited more upregulated genes compared to control samples **(Fig. 5i-l).** Similarly, pan-hormonal, macrophages, endothelial cells, and stem cells all exhibited more downregulated genes in PitNETs compared to matched cell types in control samples **(Fig. 5b-h)**, consistent with reports in which benign tumor cells will downregulate genes in stromal cells to eliminate gene expression patterns from stroma that could suppress tumor growth while malignant tumor cells will upregulate genes in the stroma to cause the stroma to actively support tumor growth^68^. Intriguingly, gene ontology enrichment analysis identified a loss of protein translation at the pre-synapse, synapse, and post-synapse as most significant findings across all normal monohormonal cells in tumor samples **(Fig. 5b-h, Supplemental Table s6),** indicating these cells are compromised in maintaining cellular connections, unlike in control samples where such networks are typically intact and functional **(Fig. 4d)**, suggesting that the loss of cell cohesiveness that neuropathologists describe as a defining characteristic of PitNETs^69^ is not just found in PitNET tumor cells but in the normal glandular cells in close proximity to them. Additionally, macrophages in tumor samples exhibited a general loss of regulation of immune system processes and leukocytic activation **(Fig. 5e, Supplementary Table 7)**, suggesting that tumor-associated macrophages in PitNETs are in an immunosuppressed state.

**Figure 5:**
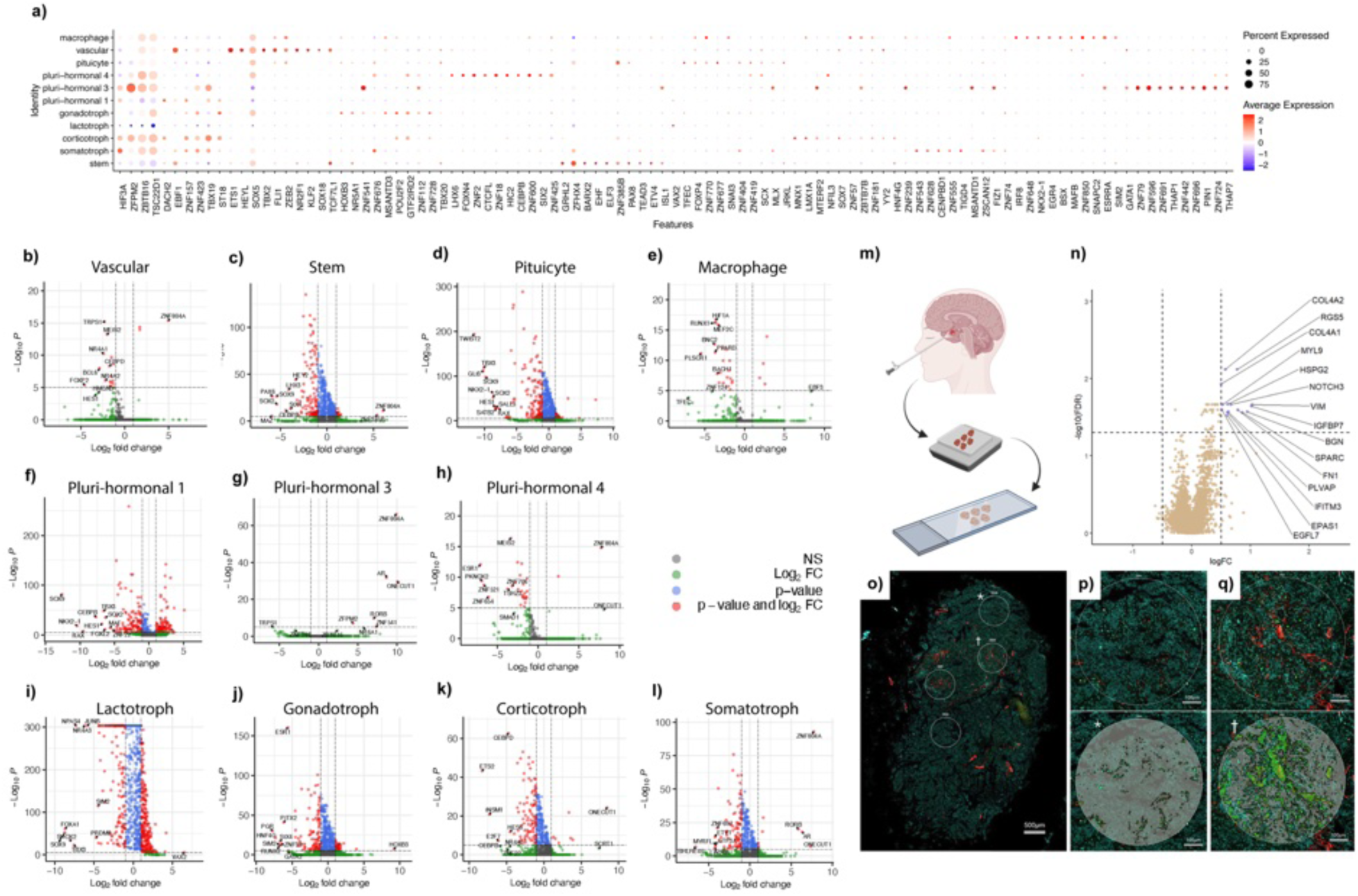
Transcriptomic analysis of the pituitary microenvironment in PitNETs compared to normal control pituitaries. (**a**) transcriptomic expression of DEGs of stromal nuclei in PitNETs relative to their counterpart nuclei in normal pituitary controls. (**b-l**) Volcano plots of all DEGs associated with each stromal cell identity, plotting the expression level of each gene in tumor stroma relative to normal pituitary stroma. (**m**) workflow of spatial transcriptomic analysis of tumor samples. Frozen pituitary adenomas from multiple patients were embedded into a single paraffin block after thawing and fixation, then mounted on a slide for GeoMx spatial transcriptomics. (**n**) Differential expression analysis comparing areas with high vascularization against those with low or no vascularization highlight 15 DEGs positively enriched for in vascularized areas. (**o**) An example of a tumor with four of the GeoMx ROI’s, represented as white circles. ⍺SMA is stained for in red with Hoechst 33342 in teal and CD68 in green. P) The top ROI in panel C representing an area of low vascularization with and without the GeoMx segmentation for tumor segments. The tumor segment (grey) excludes ⍺SMA positive pixels and CD68 positive pixels. Q) The ROI second from the top in panel C, representing an area of high vascularization with and without the GeoMx segmentation. The collected transcripts analyzed here are only from the tumor component, excluding the ⍺SMA segment, which is highlighted in green. DEG: differentially expressed gene; NS: not significant; FC: fold-change.

To characterize the vascular microenvironment of PitNETs, we identified areas of high and low vasculature density in PitNET paraffin slides and conducted spatial transcriptomic analysis **(Fig. 5m-q).** This revealed 15 upregulated DEGs in PitNET cells surrounding highly vascularized areas compared to those in low vascularized regions **(Fig. 5n),** including genes *COL4A2*, *RGS5*, *COL4A1*, *MYL9*, *HSPG2*, *NOTCH3*, *VIM*, *IGFBP7*, *BGN*, *SPARC*, *FN1*, *PLVAP*, *IFITM3*, *EPAS1*, and *EGFL7*. Many of these genes are associated with pro-angiogenic and tumorigenic activities^70–73^. Notably, *VIM* enhances tumor malignancy through its role in epithelial-mesenchymal transition (EMT) and is a target for anti-angiogenic immunotherapy^74^. Our findings suggest that crosstalk between PitNET cells and their local vasculature may mediate specific transcriptomic alterations in PitNET cells, making these cells more tumorigenic than those that do not interact with vessels, capturing a spatial-dependent heterogeneity within PitNETs beyond the interactions identified from scRNA or snRNA data.

## 3. Discussion

Pituitary neuroendocrine tumors (PitNETs) represent a unique convergence of developmental biology and oncogenesis, where the gland’s inherent plasticity is co-opted to drive tumor heterogeneity. Leveraging single-nuclei RNA sequencing (snRNA-seq) and spatial transcriptomics, we decode this complexity, revealing PitNETs not as monolithic entities but as dynamic ecosystems shaped by lineage fluidity and microenvironmental crosstalk. This plasticity-driven heterogeneity challenges the current WHO classification framework—anchored in static lineage markers such as PIT1, TPIT, and SF1—and repositions these tumors along a transcriptomic continuum where lineage boundaries are fluid and context-dependent, necessitating a revised diagnostic paradigm.

Central to our findings is the identification of plurihormonal neuroendocrine cells in both normal and neoplastic pituitary tissues. The persistence of these cells in PitNETs suggests that developmental plasticity is not merely a vestige of normal differentiation but a possible driver of tumorigenesis. In the normal adult pituitary gland, such plasticity likely facilitates adaptive responses to physiological demands, such as lactation or stress. In tumors, however, these cells—which originate early in pseudotime trajectories, retain stem-like transcriptional programs, and acquire hormonal signatures—are co-opted to sustain transcriptional diversity. These unique characteristics may enable tumor cells to evade lineage-targeted therapies and tumor suppression mechanisms.

Our trajectory analysis reveals two distinct oncogenic pathways in PitNETs: stem cell-derived tumors (e.g., somatotroph and null cell adenomas) and differentiated-cell-derived tumors (e.g., prolactinomas and corticotroph adenomas). Strikingly, pituitary stem-derived tumors exhibit fewer somatic copy-number alterations (CNAs) despite their primitive state—a paradox when contrasted with malignancies like gliomas, where cancer stem cells drive hypermutant aggression^75^. This dichotomy suggests that, in contrast to the genomic instability of cancer stem cells which have likely lost some capacity for mutational correction, pituitary stem cell-derived PitNETs may retain the developmental checkpoints found within pituitary stem cells^76^ to balance proliferation and differentiation, mirroring the genomic stability of fetal pituitary progenitors. In contrast, differentiated-cell-derived tumors may accumulate CNAs under chronic secretory stress, reflecting a trade-off between hormonal hyperactivity and genomic fidelity. These observations hint at a broader principle: PitNETs may occupy a middle ground between benign self-limited growth and aggressive transformation, offering a model to study how tumors evolve within constrained microenvironments.

PitNETs co-opt the pituitary’s native communication networks, repurposing key signaling pathways to promote tumor growth and survival. For instance, Ephrin signaling, essential for pulsatile hormone secretion in the healthy pituitary, may be hijacked to synchronize proliferation and immune evasion. Our CellChat analysis also identifies conserved ligand-receptor interactions, such as paracrine growth factor NRG1-ERBB4, which play roles in synaptic plasticity and cytoarchitecture remodeling^77^. By mimicking the normal pituitary’s intrinsic ability to coordinate complex cellular functions, these tumors may form self-organized collectives that evade immune responses and resist single-target therapies, emphasizing the need for therapeutic approaches that target broader signaling networks.

Furthermore, spatial transcriptomics localizes PitNET tumor cell-stroma crosstalk to perivascular niches. There is converging evidence that the tumor vasculature may function as a central hub to promote cancer stemness, and thereby stimulate tumor growth independently of its role as a supplier of oxygen and nutrients^78^. This observation has given rise to the term perivascular niche which refers to a microenvironment generated by endothelial cells and mural cells that surround and support blood vessels, along with nonvascular cells such as fibroblasts. In a variety of cancers, the perivascular niche affects the behavior of adjacent tumor cells, including by promoting their stemness. In the case of PitNETs, spatial transcriptomics revealed that tumor cells in the perivascular niche upregulate pro-angiogenic factors and EMT inducers, reflecting the bidirectional signaling that occurs between tumor cells and vascular cells in the perivascular niche. Tumor cells within these perivascular niches also upregulate matrix proteins (i.e., COL4A1, FN1), generating a provisional vessel-adjacent scaffold that could foster perivascular invasion—a mechanism previously described in metastatic carcinomas^79,80^ but less well-documented in benign neoplasms. Together, these findings reveal how PitNETs exploit both developmental signaling and stromal remodeling.

While our study provides important insights into PitNET biology, several limitations warrant consideration. First, despite analyzing the largest snRNA-seq dataset to date, rare PitNET subtypes (e.g., thyrotroph tumors) remain underrepresented, potentially obscuring subtype-specific mechanisms. Second, our findings implicating developmental plasticity and stromal crosstalk in tumorigenesis are primarily correlative; functional validation in *in vivo* models is needed to establish causality. Finally, our use of transcriptomic data may overlook post-transcriptional regulatory mechanisms, including alternative splicing, which could provide deeper insights into tumor heterogeneity^7^. Addressing these limitations in future studies will refine our understanding of PitNET heterogeneity and accelerate therapeutic translation.

Our work redefines PitNETs as ecosystems shaped by developmental plasticity and microenvironmental hijacking. As a gland central to physiological homeostasis, it is fitting that its tumors exhibit a biology as adaptable and interdependent as the systems they disrupt. These insights may extend beyond PitNETs, offering a framework to investigate lineage fluidity and niche dependencies in other endocrine neoplasms where similar mechanisms may take place.

## 4. Methods

### Human Tissue Procurement

Pituitary tumor samples (*n* = 11) were obtained from patients who underwent transsphenoidal endoscopic surgery at the University of California, San Francisco (UCSF) Department of Neurosurgery between 2022 and 2023. Non-tumor pituitary samples (*n* = 2) were acquired from UCSF Neurodegenerative Disease Brain Bank autopsies in 2023 with a postmortem interval of less than 10 hours. Final neuropathology diagnoses were 1) no disease and 2) Alzheimer’s disease. All tissues were snap-frozen in -80 degrees Celsius immediately upon collection. Tumor samples were histologically stained and pathologically diagnosed using standard WHO criteria. All procedures were approved by the UCSF Institutional Review Board (IRB) for human tissue research.

### Nuclei Isolation

For snRNA-seq, nuclei were extracted from frozen tissues using the ‘Van Helsing’ protocol (Martelotto, L., Centre for Cancer Research, Victorian Comprehensive Cancer Centre; protocol available at 10x Genomics dx.doi.org/10.17504/protocols.io.bdeni3de). Frozen tissues were mechanically dissociated in 500 µL lysis buffer (Nuclei EZ Lysis Buffer, MilliporeSigma) using a Dounce homogenizer. Lysates were filtered through a 30-µm strainer, pelleted, washed in nuclei wash buffer (10x Genomics), and stained with Vybrant DyeCycle Ruby Stain (Fisher Scientific) for sorting on a Sony SH800 sorter.

### SnRNA cDNA Library Generation

Isolated nuclei were resuspended in cold DPBS containing 1% BSA and 0.5 U/µL RNase inhibitor (Thomas Scientific, #C756R82). After sorting, 15,000 nuclei per sample were loaded into 10x Genomics capture reactions. Libraries were prepared using the Chromium Next GEM Single Cell 3ʹ Reagent Kit v3.1 (Dual Index, 10x Genomics, #CG000315) following the manufacturer’s protocol. Sequencing was performed on an Illumina NovaSeq with recommended parameters (28 × 10 × 10 × 90 cycles).

### SnRNA Data Processing

Raw sequencing data were processed using 10x Genomics’ Cell Ranger Count (v7.0.1) on the 10x Genomics Cloud Analysis platform. Cell/feature matrices were imported into RStudio and converted to Seurat objects (v4.2.0). Low-quality cells and doublets were filtered by excluding outliers with extreme gene counts or mitochondrial contamination (>5%). After preprocessing, mitochondrial RNA accounted for 0.25% ± 0.71% (mean ± SD) of nuclear RNA. Data were log-normalized, scaled, and integrated using canonical correlation analysis (CCA) to mitigate batch effects. Clustering resolutions (0–1.5, increments of 0.1) were optimized to maximize cluster distinctiveness while minimizing nucleus reassignment.

### Cluster Annotation

Cluster annotation was performed via consensus across three independent methods to ensure labeling accuracy. First, label transfer was conducted using the SingleR package, referencing both the standard human atlas and the Darmanis brain atlas. Second, canonical pituitary cell markers were used to compute module scores for each cluster. Finally, tumor nuclei were co-embedded with a normal pituitary snRNA-seq atlas (derived from two control samples) to validate cluster identities. Annotations were assigned based on concordance across methods and the confidence scores of each labeling approach.

### Pseudobulk and Copy Number Variation (CNV) Analysis

Pseudobulk transcriptomes were hierarchically clustered by Euclidean distance. CNVs were inferred using InferCNV (v1.14.2), comparing tumor nuclei to non-tumor controls. Tumors were reclustered by CNV magnitude and genomic location.

### Differential Gene Expression and Ontology Analysis

Differentially expressed genes (DEGs) were identified using Seurat’s FindMarkers (Wilcoxon test, Bonferroni-adjusted *p* < 0.05). Results were visualized via DotPlots (Seurat) and EnhancedVolcano (v1.16.0). Gene ontology was performed using ToppGene.

### Trajectory Analysis

Monocle (v2.26.0) reconstructed pseudotime trajectories for stem/neuroendocrine nuclei. Stem cells were designated as root notes. Pseudotime-correlated DEGs were cross-referenced with cluster markers, and ontology analysis identified pathways linked to tumorigenesis. Prolactinomas were stratified into early/late pseudotime states for phase-specific DEG analysis.

### Cell-Cell Communication Analysis

CellChat (v1.6.1) inferred ligand-receptor interactions using the reference CellChatDB.human. Pathway activity was quantified by ligand-receptor co-expression, and cluster-specific networks were computed. Signaling pathways were clustered by interaction similarity, and cell clusters were meta-clustered based on communication patterns.

### Spatial Transcriptomics Tissue Preparation and Staining

Frozen pituitary adenomas were thawed, fixed in 4% paraformaldehyde, paraffin-embedded, and sectioned at 5 µm thickness. Deparaffinized slides underwent spatial whole-transcriptome profiling on the GeoMx Digital Spatial Profiler (DSP)(Bruker NanoStrings). Antigen retrieval (20 minutes) and Proteinase K digestion (0.1 mg/mL, 15 minutes) preceded staining with AF532-conjugated anti-CD68 (1:100, Novus, #NBP2-34587AF532), AF647-conjugated anti-α-smooth muscle actin (αSMA; 1:200, Novus, #NBP2-3522AF647), and SYTO13 nuclear dye (1:10, Bruker).

### Spatial Transcriptomics Imaging and Analysis

Morphology marker images were acquired using FITC (30 ms exposure), Cy3 (300 ms), and Cy5 (300 ms) fluorescence channels. Regions of interest (ROIs) were selected based αSMA+ and αSMA-expression for spatially resolved transcript capture. Raw sequencing data were normalized to the geometric mean of housekeeping gene expression levels and analyzed using GeoMx Digital Spatial Profiler (DSP) software. Differentially expressed genes (DEGs) were defined by a Benjamini–Hochberg *FDR* < 0.05 and |log2(fold change)| > 0.5.

## Acknowledgements

The authors thank the study participants at the University of California, San Francisco, for their valuable tissue donations. They also acknowledge the staff and study coordinators at the UCSF Brain Tumor Center and Neurodegenerative Disease Brain Bank. This research was supported by the following funding sources: NIH 1R01CA227136 (M.K.A.), 2R01NS079697 (M.K.A.), 1R01NS123808 (M.K.A.), UCSF Resource Allocation Program (M.K.A.), P30 AG062422 (L.T.G.), K24 AG053435 (L.T.G.), and the Milky Way Research Foundation (P.G.)

## 5. Contributions

R.C.O., J.Y.O., J.J. contributed equally as co-first authors. R.C.O., J.Y.O., J.J. and M.K.A. designed the study. R.C.O., J.Y.O., J.J. performed single-nuclei RNA-sequencing experiments and data analysis with significant contributions from A.S., M.L., and H.K.. A.J.E., N.B., and P.G. performed spatial transcriptomics experiments and data analysis. W.W.S., L.T.G., and A.D. contributed to critical data collection and interpretation. R.C.O., J.Y.O., J.J. wrote the manuscript, and all authors contributed to the writing and provided comments. M.K.A. provided supervision.

---

**Supplementary Figure 1:**
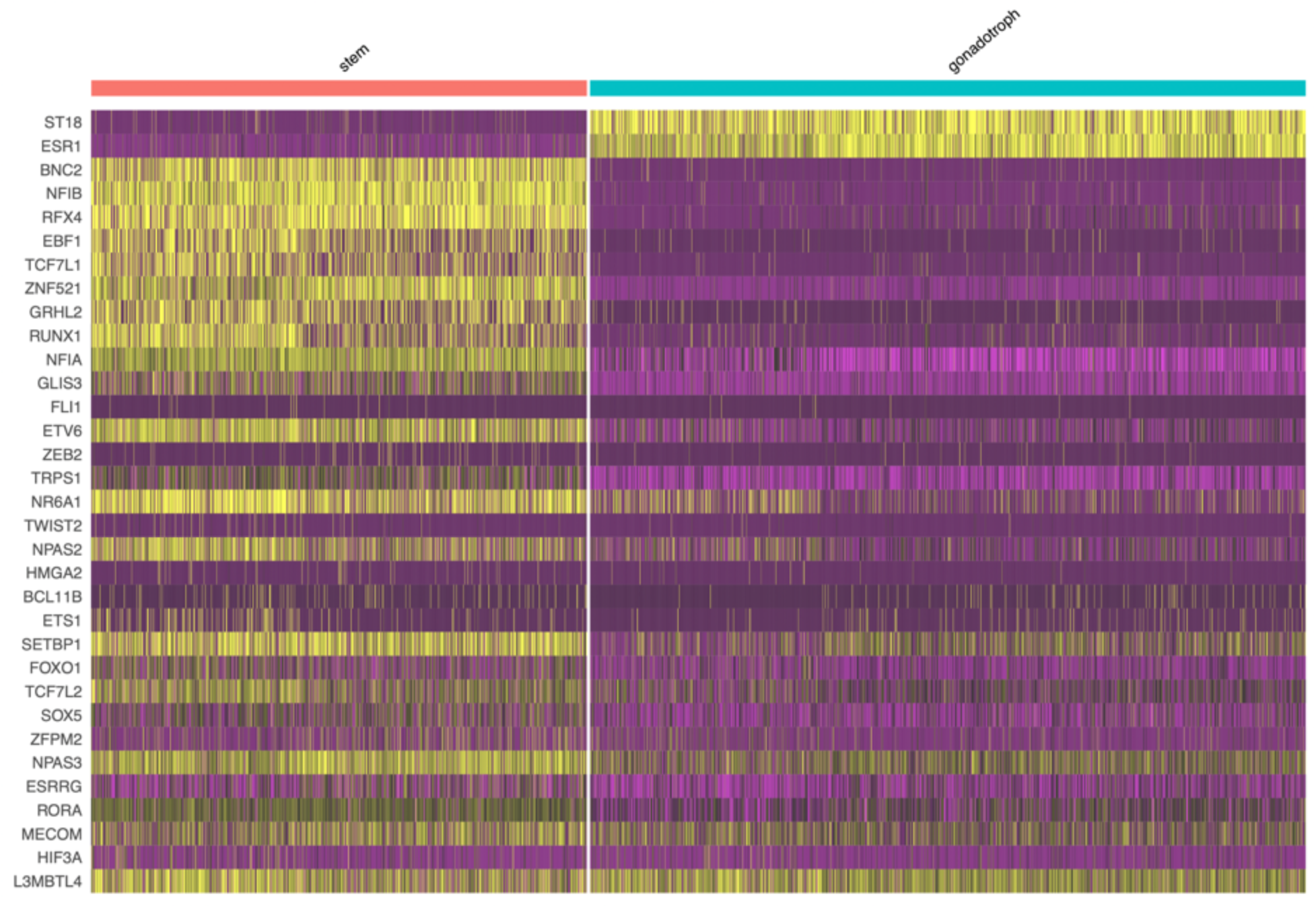
Transcriptomic heatmaps of pseudotime-correlated differentially expressed genes (DEGs) in normal gonadotroph cells. DEGs expression is plotted with respect to trajectory-analysis pseudotime and visualized in stem cells vs normal gonadotroph cells.

**Supplementary Figure 2:**
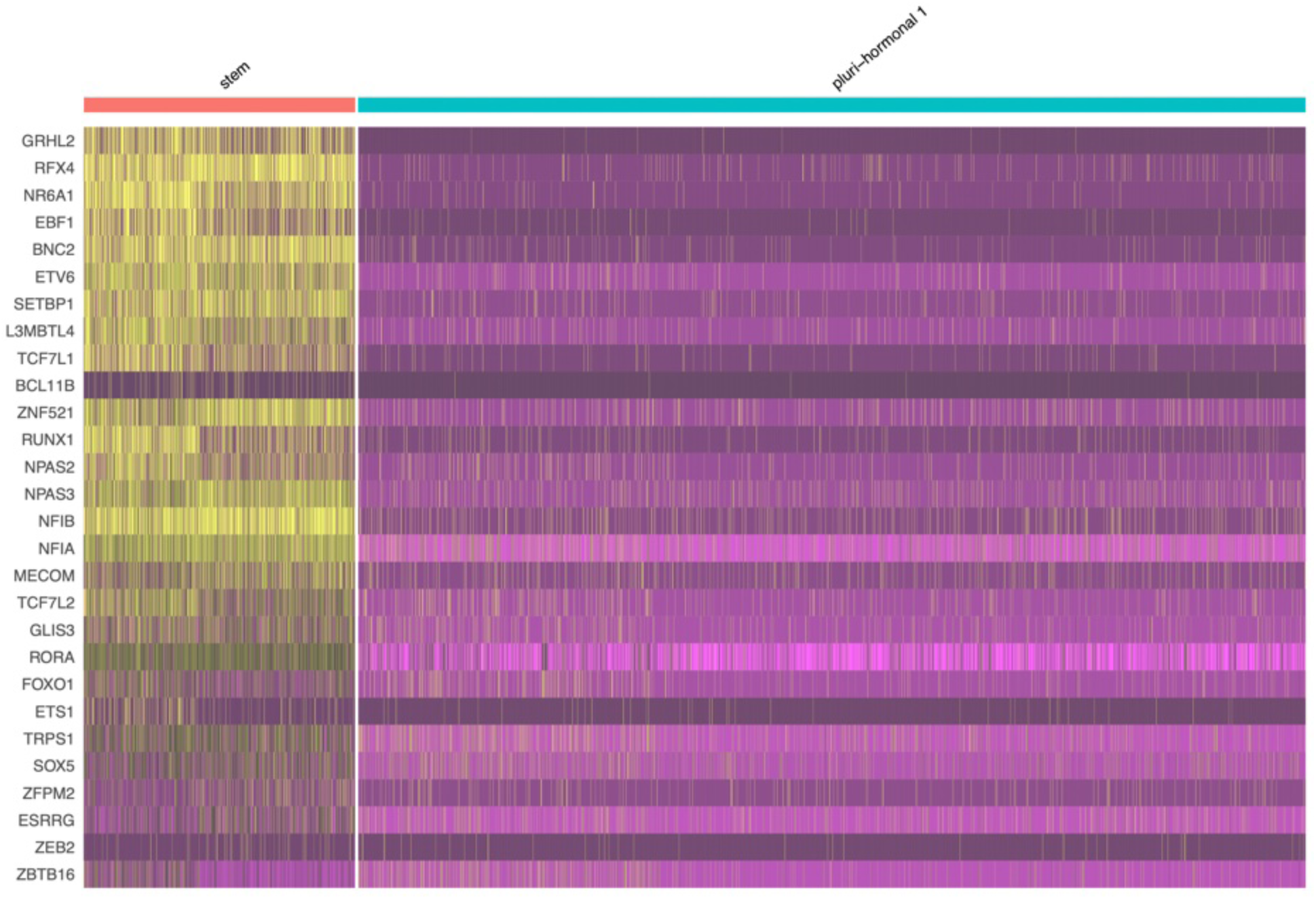
Transcriptomic heatmaps of pseudotime-correlated differentially expressed genes (DEGs) in normal gonadotroph cells. DEGs expression is plotted with respect to trajectory-analysis pseudotime and visualized in stem cells vs cells in the “pluri-hormonal 1” cluster.

**Supplementary Figure 3:**
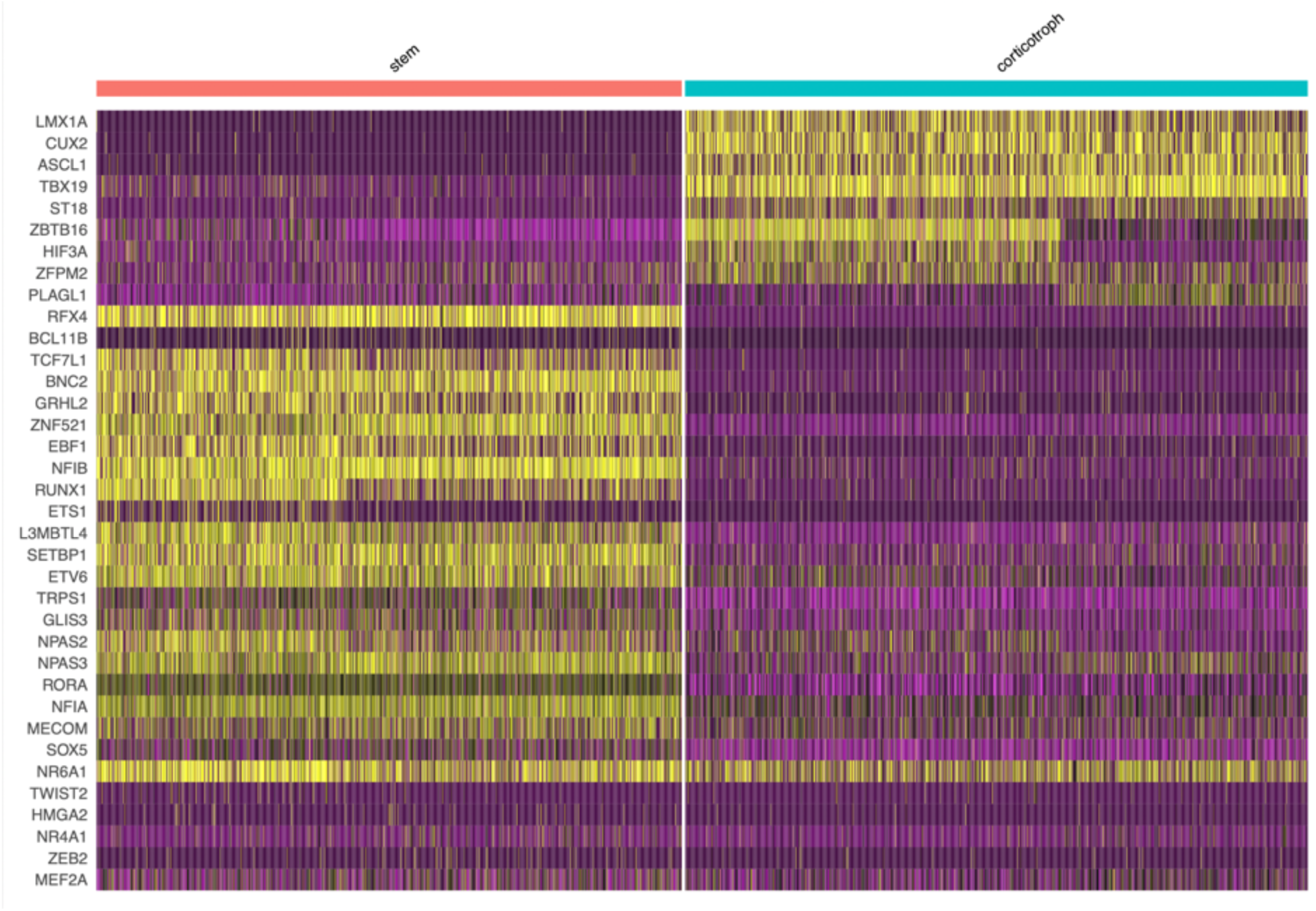
Transcriptomic heatmaps of pseudotime-correlated differentially expressed genes (DEGs) in normal gonadotroph cells. DEGs expression is plotted with respect to trajectory-analysis pseudotime and visualized in stem cells vs normal corticotroph cells.

**Supplementary Figure 4:**
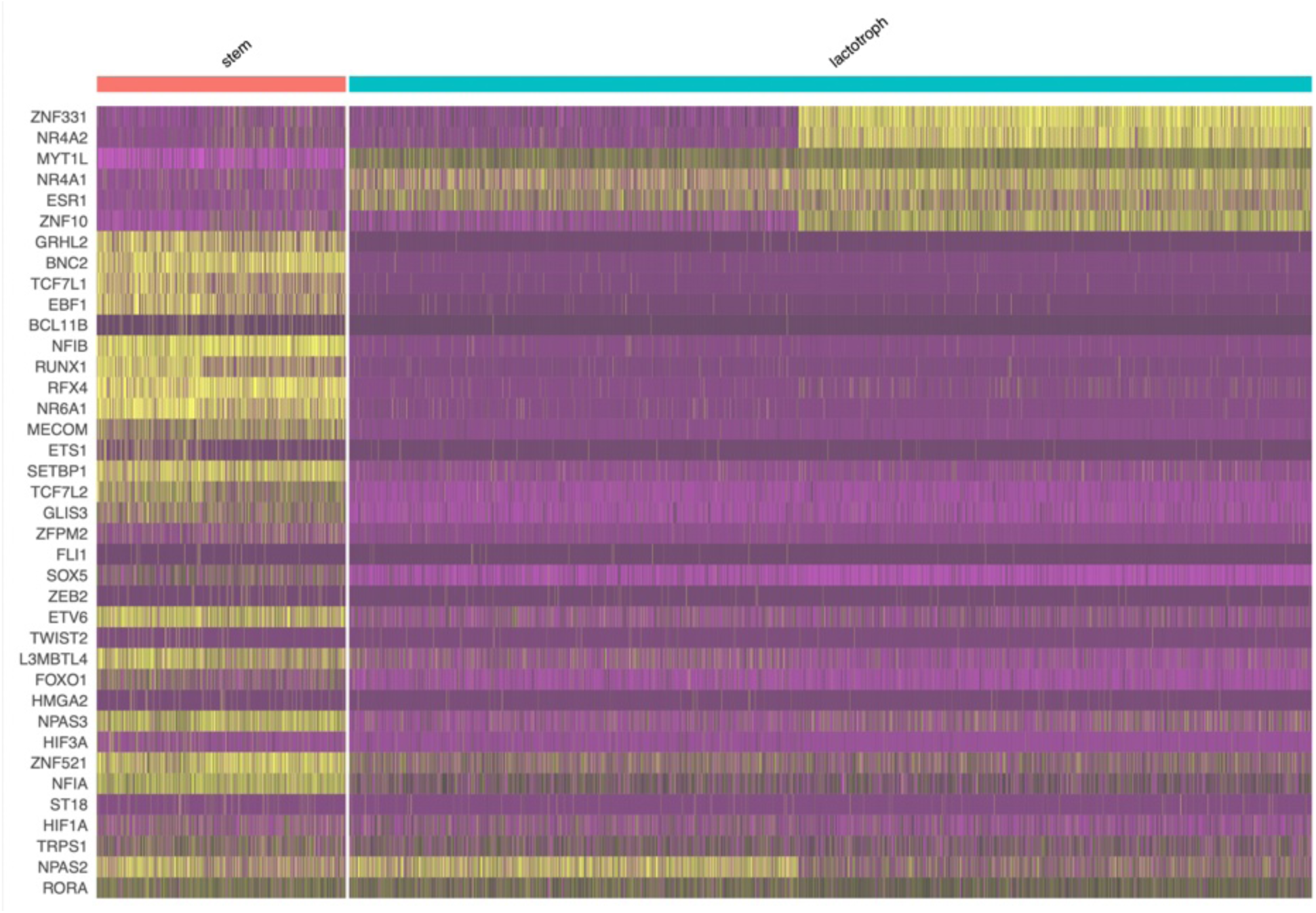
Transcriptomic heatmaps of pseudotime-correlated differentially expressed genes (DEGs) in normal gonadotroph cells. DEGs expression is plotted with respect to trajectory-analysis pseudotime and visualized in stem cells vs normal lactotroph cells.

**Supplementary Figure 5:**
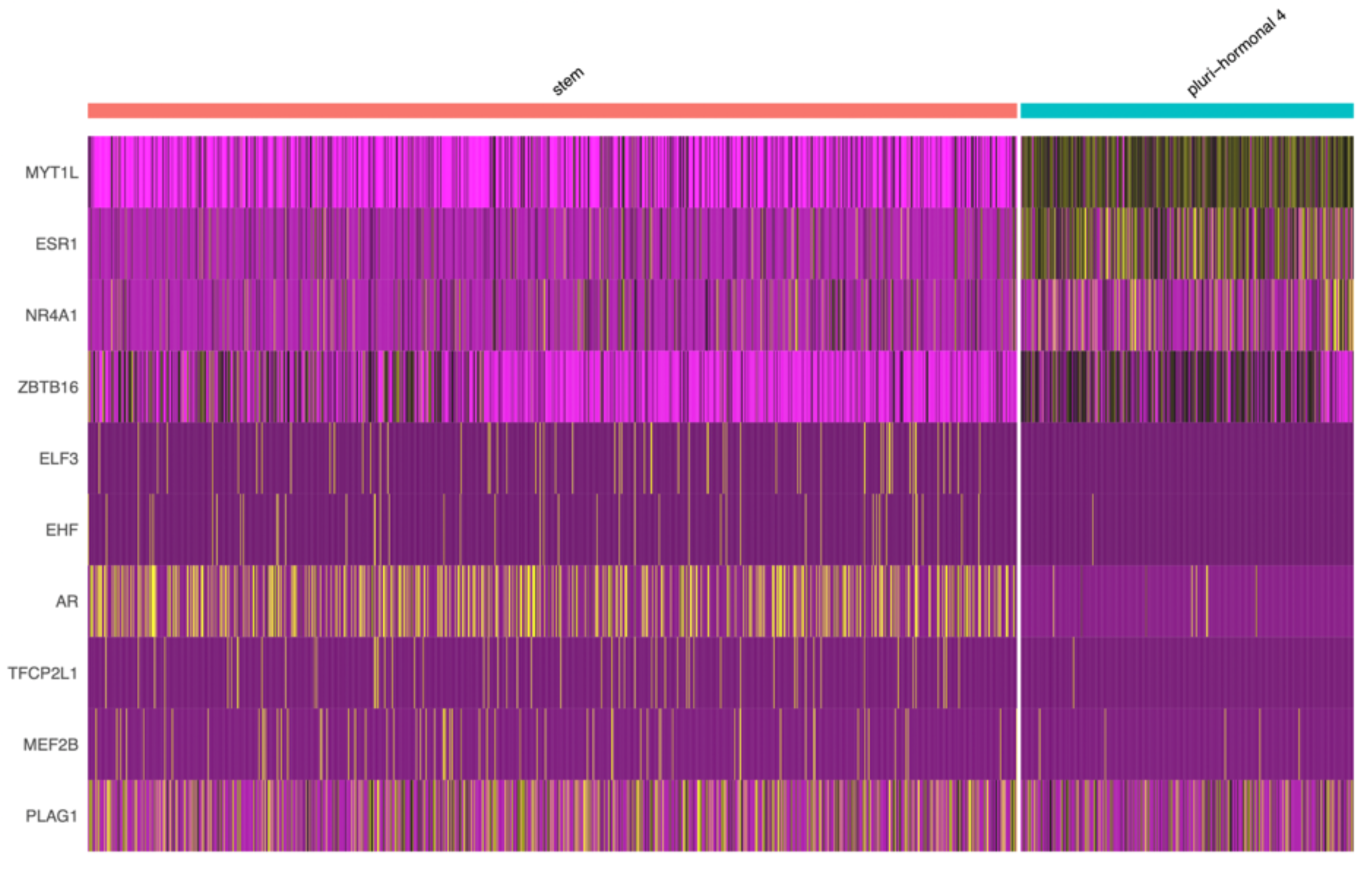
Transcriptomic heatmaps of pseudotime-correlated differentially expressed genes (DEGs) in normal gonadotroph cells. DEGs expression is plotted with respect to trajectory-analysis pseudotime and visualized in stem cells vs cells in the “pluri-hormonal 4” cluster.

**Supplementary Figure 6:**
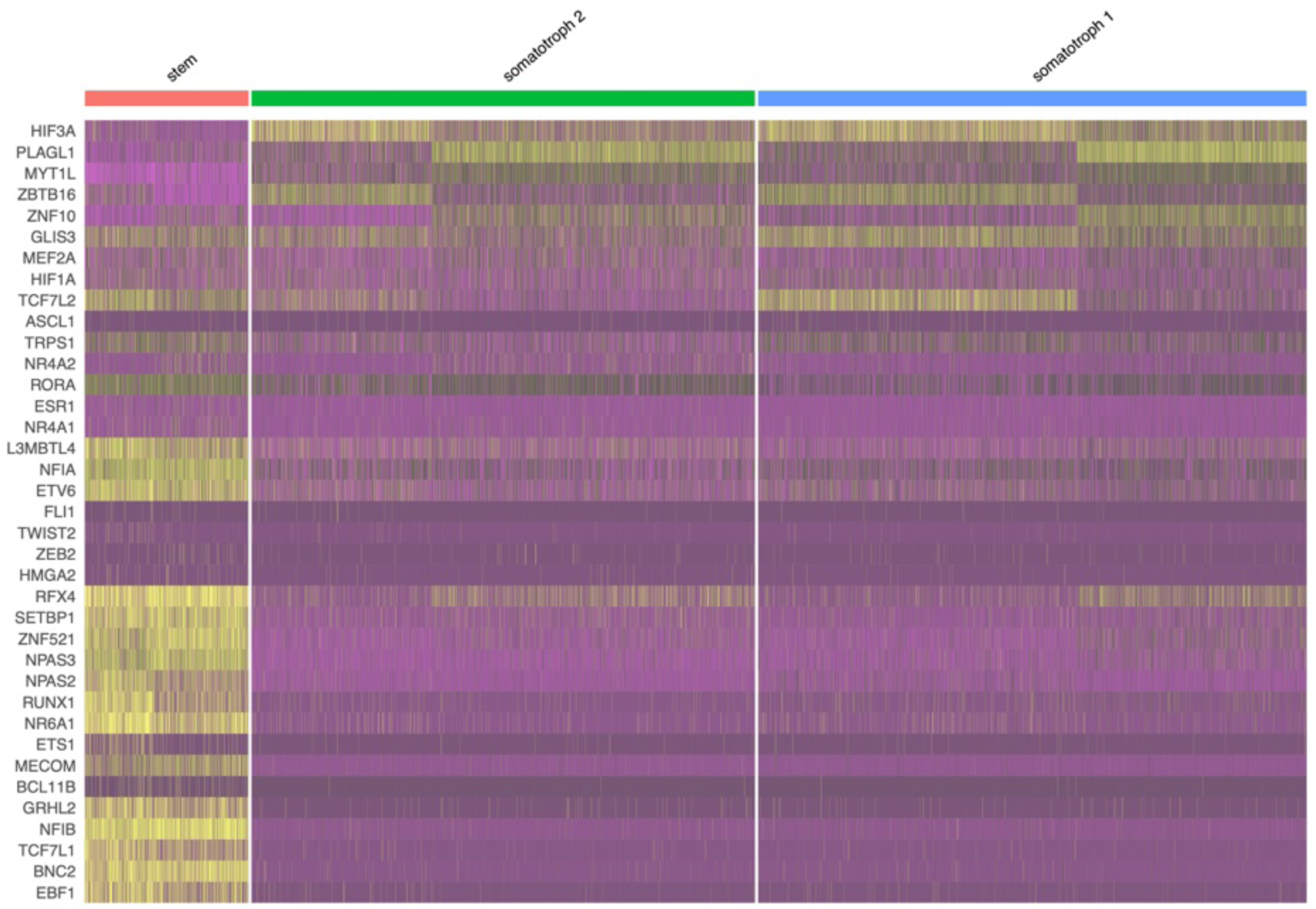
Transcriptomic heatmaps of pseudotime-correlated differentially expressed genes (DEGs) in normal gonadotroph cells. DEGs expression is plotted with respect to trajectory-analysis pseudotime and visualized in stem cells vs cells in the “somatotroph 1” and “somatotroph 2” clusters.

**Supplementary Figure 7:**
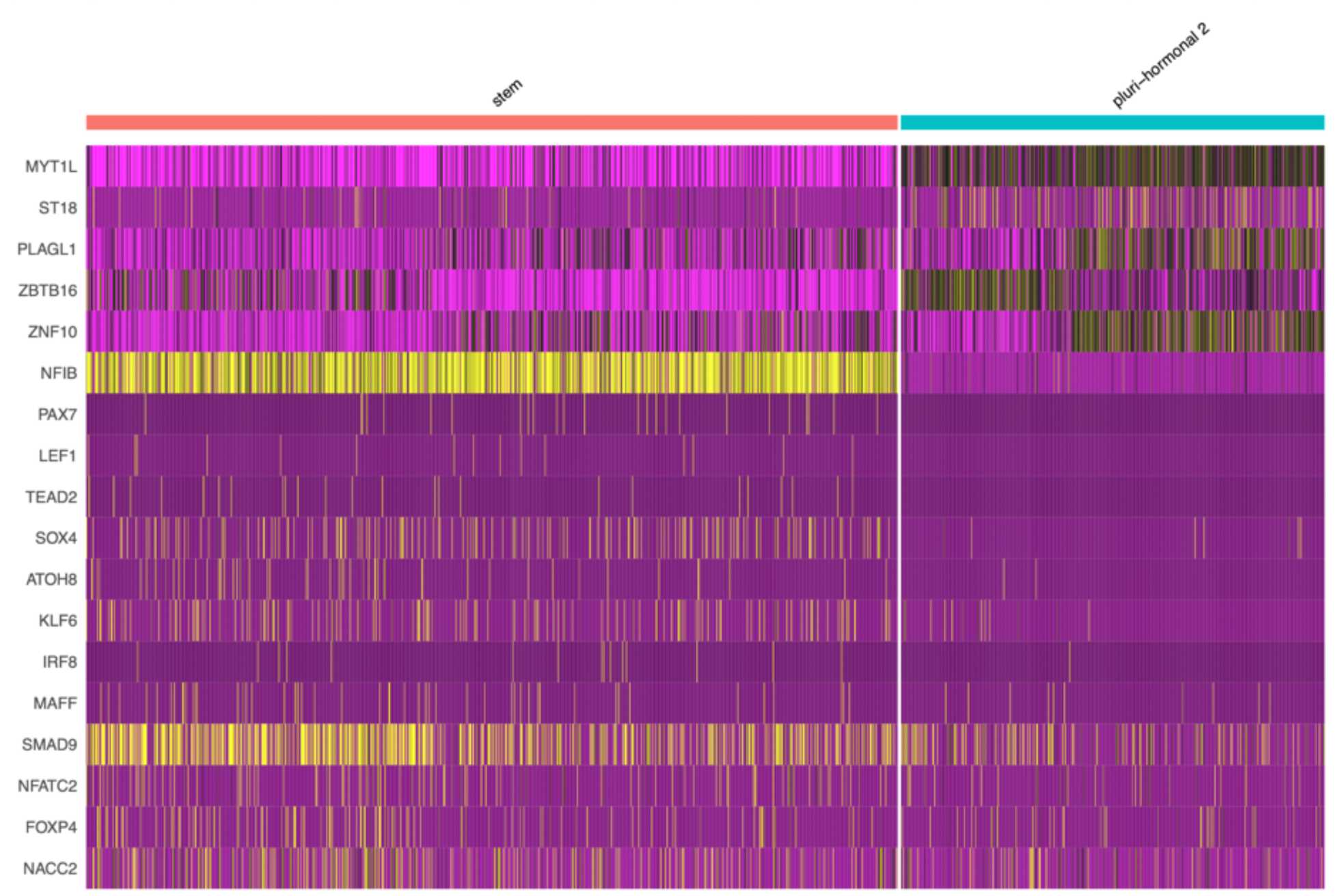
Transcriptomic heatmaps of pseudotime-correlated differentially expressed genes (DEGs) in normal gonadotroph cells. DEGs expression is plotted with respect to trajectory-analysis pseudotime and visualized in stem cells vs cells in the “pluri-hormonal 2” cluster.

**Supplementary Figure 8:**
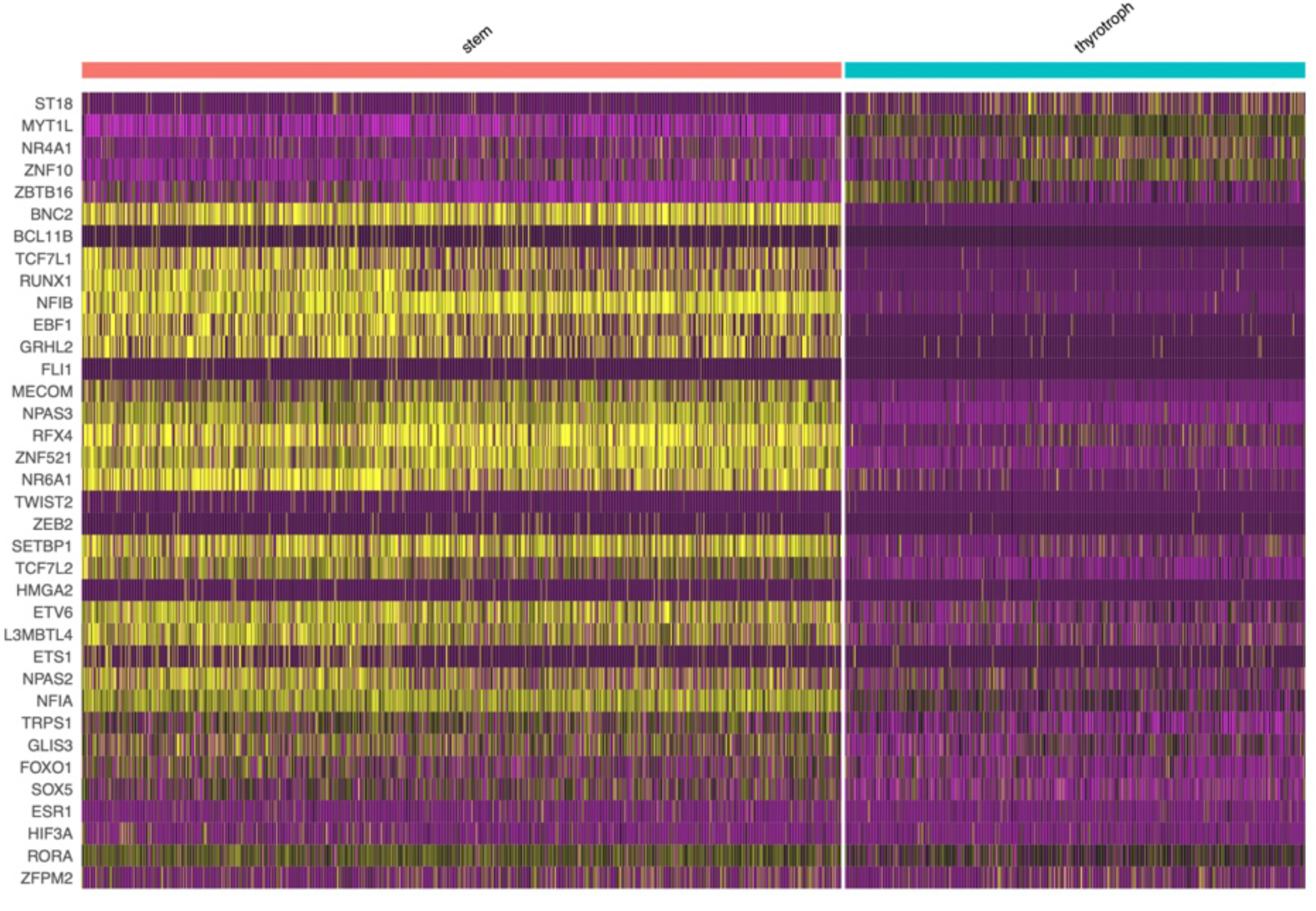
Transcriptomic heatmaps of pseudotime-correlated differentially expressed genes (DEGs) in normal gonadotroph cells. DEGs expression is plotted with respect to trajectory-analysis pseudotime and visualized in stem cells vs thyrotroph cells.

## Notes

### Competing Interest Statement

The authors have declared no competing interest.

### Summary of Updates

We have included additional authors who have contributed to the paper.

